# Extracellular matrix changes in colorectal carcinoma and correlation with lymph node metastasis

**DOI:** 10.1101/2021.01.05.425386

**Authors:** Thérèse Rachell Theodoro, Rodrigo Lorenzetti Serrano, Karine Corcione Turke, Sarhan Sydney Saad, Marcelo Augusto Fontenelle Ribeiro Junior, Jaques Waisberg, Maria Aparecida Silva Pinhal

## Abstract

The process of proliferation and invasion of tumor cells depends on changes in the extracellular matrix (ECM) through the activation of enzymes and alterations in the profile of ECM components. We aimed to investigate the mRNA and protein expression of ECM components such as heparanase (HPSE), heparanase-2 (HPSE2), matrix metalloproteinase-9 (MMP-9), and syndecan-1 (SYND1) in neoplastic and non-neoplastic tissues of patients with colorectal carcinoma (CRC). It is a cross-sectional study in which twenty-four adult patients that had CRC were submitted to resection surgery. We analyzed the expression of HPSE, HPSE2, MMP-9, and SYND1 by quantitative RT-PCR and immunohistochemistry. Differing from most of the studies that compare the mRNA expression between tumor samples and non-neoplastic tissues, we decided to investigate whether variations exist in the expression of the ECM components between the affected tissue and nontumoral tissue collected from the same patient with CRC. We removed both tissue samples immediately after the surgical resection of CRC. The data showed higher mRNA and protein expression of HPSE2 (P = 0.0058), MMP-9 (P = 0.0268), and SYND1 (P = 0.0002) in tumor samples compared to the non-neoplastic tissues, while there was only an increase in the level of HPSE protein in tumor tissues. A greater expression of HPSE2 was observed in patients with lymph node metastasis (P = 0.048), suggesting that such protein can be a marker of lymph node metastasis in CRC.

## Introduction

Colorectal carcinoma (CRC) is the third-leading source of death from cancer worldwide, and the second-most frequent cause of death in developed countries [1]. Despite being the primary procedure in the treatment of CRC, surgical resection has limited efficacy when the neoplasia presents local invasion or distant metastases [2]. It is essential to understand the mechanisms that allow the disease’s progression and find markers that indicate the prognosis of CRC [3].

Concerning the mechanism of neoplastic invasion and metastasis production process in solid tumors, great importance has been attributed to the interactions between tumor cells and the stroma surrounding the neoplasia [4,5].

Cell proliferation and the ability of neoplastic cells to invade tissues are dependent on the loss of the cell-cell interaction process that occurs in the extracellular matrix (ECM). This loss leads to the degradation of ECM components such as proteins, proteoglycans, and glycosaminoglycans by activation of specific enzymes associated with carcinogenesis and tumor metastasis [6]. Loss of natural epithelial adhesion and altered expression of cell adhesion molecules are associated with the development of colorectal neoplasms [5,6].

Heparan sulfate (HS) proteoglycans are cell membrane and ECM molecules, and they act as a growth factor and cytokine coreceptors involved with cellular signaling [3,6]. Cell surface HS proteoglycans act as regulators of cell migration and proliferation [4].

Syndecan-1 (SYND1, CD138), a member of the syndecan family of heparan sulfate (HS) proteoglycans, is encoded by the *SYND1* gene located on chromosome 2p24.15 [6]. SYND1 is predominantly expressed in the basolateral membrane of epithelial cells [2,4,5]. The binding of SYND1 with cytokine molecules and EMC components is carried out through the HS chain [4,5], which allows SYND1 to play an important role in the remodeling of EMC, in addition to participating in physiological functions such as tissue repair, regulation of immune function, and controlling inflammation [5, 6]. SYND1 is involved in molecular pathways during carcinogenesis related to cell proliferation, apoptosis, angiogenesis, and tumor invasion [3,4]. An epithelial/mesenchymal transition event and transformed phenotype are associated with loss of SYND-1 expression, whereas this molecule is highly expressed in normal epithelial cells [3].

Heparanase (HPSE) is an endoglucuronidase that cleaves HS chains of proteoglycans at specific intrachain points [7,8]. HPSE releases oligosaccharides that bind to growth factors and cytokines and intensify the activity of such components^7^. The MEC network remodeling facilitates cell motility, angiogenesis, inflammation, coagulation, stimulation of autophagy, and exosomes production [9].

HPSE has two isoforms. The human *HPSE* gene is located on chromosome 4q21.23.23. It encodes heparanase-1 isoform (HPSE), whose action disassembles the subendothelial basement membrane and facilitates the installation of metastatic cells disseminated by the bloodstream in the tissues by cleavage of the proteoglycans’ HS [8,10]. In normal tissue and under physiological conditions, HPSE expression levels are low, and the protein is found in keratinocytes, trophoblasts, and platelets, as well as mast cells and leukocytes [11]. HPSE also stimulates tumor angiogenesis and vascularization [8,10] and is involved in cell surface HS and ECM degradation in both neoplastic and nonneoplastic tissues. The Heparanase-2 (*HPSE2*) gene is located on chromosome 10q24.2 [10] and encodes the heparanase-2 isoform (HPSE2), a cell membrane protein. HPSE2 has no enzymatic activity and appears to regulate HPSE activity [10]. The level of expression of HPSE2 is elevated in the brain, small intestine, breasts, uterus, bladder, prostate, and testis [12].

HPSE activity participates in disseminating neoplastic metastasis, and neoplasms that exhibit high HPSE expression and activity also correlate with a more aggressive tumor phenotype [7,8]. Researchers have emphasized the involvement of HPSE in exosome formation, immune system activation, autophagy, and chemo-resistance, which further the tumor response to host defense factors, which are influenced by the mediation of interference between tumor cells and tumor microenvironment [7–9]. HPSE has a protumorigenic effect by facilitating cell invasion and triggering the tumor microenvironment’s installation mainly due to its degradation activity of HS [8,13].

HPSE acts on ECM related to epithelial and endothelial cells, altering their structural integrity by cleavage of HS side chains [14–16]. This action facilitates the associated processes of neoplastic cell invasion, metastasis, and tumor angiogenesis [14,16,17]. In the overexpression of the *HPSE* gene, research has observed the spread of neoplastic cells in the lungs, liver, and bones, which together with the significant decrease in the metastatic potential of neoplastic cells caused by silencing the *HPSE* gene reinforces the role of HPSE in the metastasis process [17].

The removal of HS from the cell surface with bacterial heparitinase increased SYND1 shedding [14]. This result indicates that enzymatic removal or reduction in the size of HS chains influences the action of HPSE on SYND1 expression by neoplastic cells [14]. Deactivating the HPSE molecule has become an attractive therapeutic target due to the lack of identification of other molecules that perform the same function as HPSE [15,16]. Researchers have observed that HPSE has prometastatic and proangiogenic activity in primary malignant neoplasms [14,15,18]. Increased HPSE levels were also found to correlate with the presence of lymph nodes compromised by neoplasia and distant metastases, as well as high microvascular density and reduced survival index [16,19].

Expression of mRNA and HPSE-related proteins was observed to be markedly elevated in CRC [18,20]. Elevated levels of HPSE were related to more advanced TNM classification, vascular and lymphatic infiltration, and worsening survival because HPSE participates in tumor development and growth and facilitates metastasis implantation [16,18,20]. Research has been suggested that *HPSE2* is a tumor suppressor gene [2,10,13]. SYND1 shedding may explain HPSE2’s overexpression in CRC tissue compared to nonneoplastic tissues because HPSE2 has no enzymatic activity [10].

Matrix metalloproteinases (MMP) are calcium-dependent zinc endopeptidases [21,22] and are physiologically involved in the degradation of ECM components, embryogenesis, reproduction, angiogenesis, bone development, wound healing, and cell migration [23]. Matrix metallopeptidase 9 (MMP-9) has its genetic locus on chromosome 20q11.2-q13.1 and is produced by epithelial cells, macrophages, granulocytes, T-cells, keratinocytes, dendritic cells, osteoblasts, and fibroblasts [24–26].

MMP is synthesized by tumor cells and stromal cells surrounding the neoplasia and can degrade the basal membranes, often the first step of carcinoma invasion. MMP facilitates tumor progression, including invasion, metastasis, growth, matrix remodeling, and angiogenesis, by increasing the concentration of vascular endothelial growth factor (VEGF) [22,23]. Metalloproteinase-9 (MMP-9) is also active in the formation of early metastatic niches [24]. Due to these activities, MMP-9 represents an important proteolytic enzyme for breaking and reconstructing ECM that influences the invasion process and the development of metastases in CRC evolution [23,25,27]. In the treatment of CRC, a decrease in cancer growth and metastatic involvement was observed using of selective MMP-9 inhibitors in preclinical studies [24].

Analysis of SYND1 expression associations with pathological and prognostic aspects in patients with CRC produced controversial results concerning correlations between loss of SYND1 expression, histological grade, CRC stage, lymph nodes compromised by the neoplasia or distant metastases, TNM stage, and survival [3,28–32]. Studies have presented conflicting results regarding the role of MMP-9 expression in CRC patients’ prognoses concerning metastases, lymph nodes, and poor outcomes [33–36].

Our group previous studies showed a direct correlation between HPSE and SYND1, whereas we observed an inverse correlation between HPSE2 and HPSE and between HPSE2 and SYND1 in the colorectal adenomas [37]. We also observed an increased HPSE2 protein level and a decreased SYND1 level in CRC [2]. Therefore, knowledge of the correlation between HPSE, HPSE2, MMP-9, and SYND1, we decide to investigate the expression of such molecules in neoplastic and nonneoplastic tissues collected from CRC patients and correlate findings to the presence of lymph node metastases. Together, our findings support that HPSE2 may play a role as a marker of lymph node metastasis in CRC.

## Materials and methods

### Ethical

All procedures performed in studies involving human participants were in accordance with the ethical standards of the Institutional Research Ethics. The study was approved by the Research Ethics Committee of Federal University of São Paulo, number: 031/05, and is in accordance with the 1964 Helsinki declaration and its later amendments. The written informed consent was obtained from all patients. This article does not contain any studies with animals performed by any of the authors.

### Patients and Samples

This study was a cross-sectional study involving patients operated consecutively on CRC in 12 months period (2008 – 2009) in the Department of Gastrointestinal Surgery, State Public Servant Hospital (São Paulo, Brazil).

Immediately after resection of the large intestine, two samples were taken by a surgeon (JW) from the large bowel of each patient: one from the neoplastic region, but out from the center of the tumor, and one from a non-neoplastic area located 10 cm cranially at the border of the neoplasia. Each collected tissue was divided in three other samples: one for RNA extraction, one for immunohistochemistry analysis, and one to visualize the tissues’ anatomopathological characteristics. For RNA extraction, colorectal tissue samples were immediately placed in the RNA holder solution (BioAgency Biotechnology, São Paulo, SP, Brazil) for the preservation of samples that were subsequently submitted to extraction of total RNA for quantitative RT-PCR amplification assays, using specific oligonucleotides for each marker. Additionally, two samples, one from the neoplastic region one from the non-neoplastic area, were placed in 10% neutral buffered formaldehyde for immunohistochemistry and anatomopathological analysis.

Adult patients of both genders, without a distinction of ethnicity, were included in this study with the confirmed histological diagnosis of CRC and operated with curative intention. Patients with familial adenomatous polyposis, hereditary CRC syndromes, colorectal neoplasia other than carcinoma, inflammatory bowel disease, those submitted to neoadjuvant chemotherapy and/or radiotherapy, with synchronous/metachronous tumors elsewhere or those subjects with missing data were excluded. Overall, 50 patients were operated on curative or palliative basis for CRC, of which 26 of these patients were excluded, and 24 patients met the above criteria (Fig 1), including 12 men (50%) and 12 women (50%), all participants were white. The mean age was 67.3±9.3 (range = 40-86) years. The most relevant comorbidities were arterial hypertension (N= 14), diabetes mellitus (N= 6) and rheumatoid arthritis (N= 2). The investigation for preoperative staging was performed with abdominal CT, abdominal magnetic resonance, chest CT and determination of the serum level of carcinoembryonic antigen (CEA). Fig 1 shows the flowchart used for the selection of patients involved in the present study.

**Fig 1.**
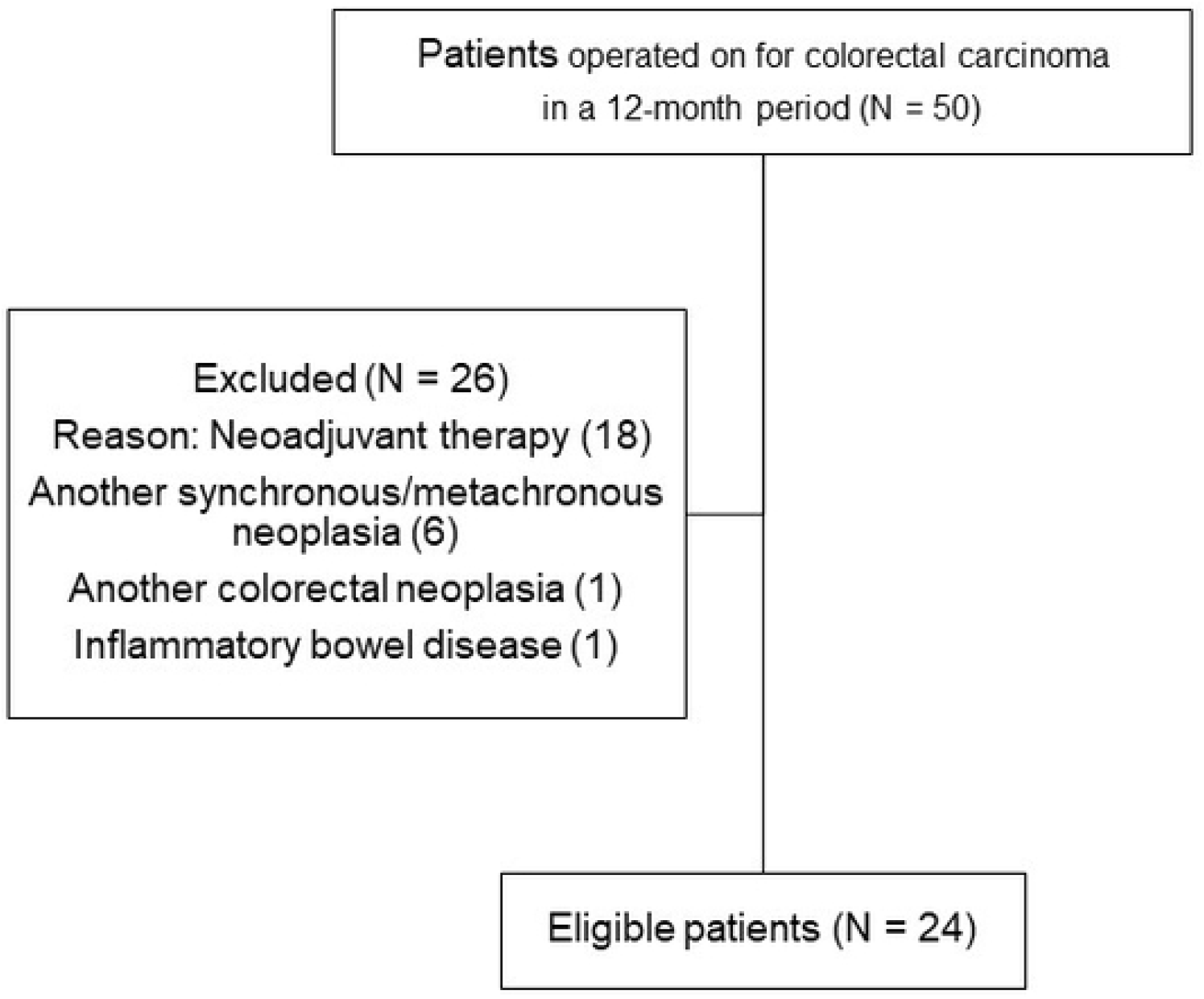
Adopted inclusion and exclusion criteria in the study. Flowchart used for selection of patients with colorectal carcinoma submitted to surgery.

The following clinical and pathological data were analyzed: gender, tumor site, tumor size, degree of tumor differentiation, wall invasion, vascular invasion (presence vs. absence), perineural invasion (presence vs. absence), and lymph nodes metastases (presence vs. absence). Only cases in which 12 or more lymph nodes were removed by the surgical procedure were included in the study.

### Total RNA Extraction

We extracted total RNA from all fractions using Trizol reagent (Ambion, Life Technologies, Foster City, CA, USA), following the manufacturer’s instructions. The integrity of the ribosomal RNA was determined by the presence of the 28S and 18S bands on the agarose gel, after electrophoresis. Total RNA from samples was quantified using NanoVeu (GE-Healthcare Limited, Buckinghamshire, UK).

### Obtaining cDNA

We performed reverse transcription using the reverse transcriptase enzyme ImPromII (Promega Co., Fitchburg, WI, USA) from 1 μg of total RNA, following the manufacturer’s instructions for obtaining the cDNA. We quantified all cDNA samples were quantified using NanoVeu (GE-Healthcare Limited, Buckinghamshire, UK), and we obtained 1 μg/μL for further real-time PCR amplification.

### Real Time RT-PCR

We used pairs of specific primers, commercially named forward and reverse primers. The expression of the HPSE mRNA isoforms (HPSE and HPSE2) was plotted against the geometric mean of the expression of endogenous reference genes, ribosomal protein 60S L13A (RPL13a), primer forward 5’TTGAGGACCTCTGTGTATTTGTCAA3’ and reverse 5’ CCTGGAGGAGAGAGAGGAAGAGA3’, and the enzyme glyceraldehyde-3 phosphate dehydrogenase (GAPDH), primers forward 5’TCGACAGTCAGCCGCATCTTCTTT3’ and reverse 5’GCCCAATACGACCAAATCCGTTGA3’, thus determining the values of (-ΔCt). WE analyzed the expression of target genes using the following primers for HPSE primers forward 5’TGGCAAGAAGGTCTGGTTAGGAGA3’ and reverse 5’GCAAAGGTGTCGGATAGCAAGGG3’; for HPSE2 primers forward 5’AGACAGAGCTGCAGGTTTGAAGGA3’ and reverse 5’AGCTTAGGAAATCGAGCCAGCCAT3’; for MMP-9 primers forward 5’GCCTGGCACATAGTAGGCCC3’ and reverse 5’CTTCCTAGCCAGCCGGCATC3’ and SYND1 primers forward 5’AGGGCTCCTGCACTTACTTGCTTA3’ and reverse 5’ATGTGCAGTCATACACTCCAGGCA3’. Trials were performed in triplicates. All primers were produced by Applied Biosystems (Carlsbad, CA, USA). We perform amplification using the Maxim SYBR Green qPCR Master Mix reagent (2X) (Applied Biosystems, Carlsbad, CA, USA), following the protocol (1.5 μM of each primer, 1 μg cDNA, 0.025 μL of the ROX 50 solution diluted 10X, and 6 μL of SYBR Green reagent 2X). We subject the mixture to a cycler for realtime amplification 7500 Real Time PCR Cycle (Applied Biosystems, Carlsbad, CA, USA), cycling at 95°C for 10 minutes, followed by 40 cycles (95°C, 15 s, 60°C, 60 s). The temperature of the dissociation or melting curve was determined for each pair of primers.

### Immunohistochemistry

The tissues were fixed in 10% formalin and embedded in paraffin, from which 5 μm sections were prepared for immunohistochemistry staining using specific antibodies as described. Anti-heparanase 1 (HPA1 C-20) polyclonal antibody (diluted 1:100); reactivity against human, mouse and rat; anti-heparanase 2 (HPA2 C-17) polyclonal antibody (diluted 1:100); reactivity against human, mouse and, rat; anti-metalloproteinase-9 (MMP-9 C20) polyclonal antibody (diluted 1:100); reactivity against mouse, rat and human, such polyclonal antibodies were obtained from Santa Cruz (Santa Cruz, CA, USA). Anti-syndecan-1 (MCA681) monoclonal antibody (diluted 1:50) reactivity against human and obtained from the Bio-Rad Company Co. (AbD Serotec^®^, Oxford, UK). The negative control of the reaction was performed in the absence of primary antibody using 1% BSA (bovine serum albumin) diluted in phosphate buffer. Immunolabeling was carried out using the avidin-biotin-peroxidase complex, following the protocol described by the manufacturer, LSAB+System-HRP kit (Dako North America, Inc., CA, USA) and 3,3’-diaminobenzidina as liquid chromogen - DAB + Substrate Chromogen System (Dako North America, Inc., CA, USA). The sections were analyzed under a TS100 Nikon Eclipse^®^ light optical microscope to identify the areas that best represented the immunolabeling. For each case, microphotographs of 640×480 pixels were obtained from consecutive and non-coinciding fields, with magnification of 10X and 40X microscope objective lenses, through an optical microscope using an Aperio CS2 (Leica Biosystems) scanner, generating digital files. The slides were then analyzed using ImageScope™ software (Aperio Technologies). All histomorphometry analysis was performed by two observers and one pathologist. A Nikon brand binocular microscope was used. Initially, an analysis of two dorsal quadrants was performed under 10Xx magnification and then the analysis was performed on 5-10 concentric fields under 40X magnification. The assessment was done using semiquantitative scoring for the intensity of immunolabeling indicated as Expression Intensity values (EIt) as described absent (zero); mild (1); moderate (2); intense (3), very intense (4).

All slides were digitalized using a TS100 Nikon Eclipse^®^ optical microscope and divided into two quadrants. For each quadrant, 10 equal and random areas were selected for a given intensity of expression, as described (zero, mild, moderate, intense and very intense). The entire procedure was performed by three different observers to avoid bias. After all quantifications were completed, they were calculated as mean and standard deviation. Both neoplastic and non-neoplastic areas were selected according to the guidance of a pathologist.

### Statistical Analysis

The qualitative variables were described by absolute and relative frequencies. For the quantitative variables, the median and the 25th, and 75th percentiles were used. The statistical tests used in the analysis were based on the type of distribution by the Kolmogorov-Smirnov test. Student’s one sample t-test for comparison of parametric independent groups and the Mann-Whitney test for nonparametric groups. The Wilcoxon test was assigned for comparison of nonparametric dependent groups. We adopted a statistical significance level of 5% (P ≤ 0.05). Two-tailed tests were applied for all statistical analyses using SPSS version 13.0 (SPSS Inc., Chicago, IL, USA) software.

## Results

No patients had distant metastases. The CRC was located in the left colon in 6 (25%) patients, right colon in 3 (12,5%) and in the rectum in 15 cases (62.5%). A right colectomy was performed in 2 (8.3%) cases, a left colectomy in 7 (29.2%), and a in 15 (62.5%). Considering all operated patients, 425 lymph nodes were removed with an average of 19.0 lymph nodes per patient. The clinical and pathological characteristics of patients operated on CRC are summarized in Table 1.

**Table 1.** Clinicopathological characteristics of patients operated on for colorectal carcinoma.

|  |  | N | % |
| --- | --- | --- | --- |
| Gender | Male | 12 | 50 |
|  | Female | 12 | 50 |
| Site | Colon | 9 | 37.5 |
|  | Rectum | 15 | 62.5 |
| Size | <5cm | 14 | 58.3 |
|  | ≥5cm | 10 | 41.7 |
| Tumor differentiation | Well-defined | 2 | 8.3 |
|  | Moderately-defined | 22 | 91.7 |
| Wall invasion | T2 | 6 | 25 |
|  | T3 | 18 | 75 |
| Vascular invasion | Yes | 8 | 33.3 |
|  | No | 16 | 66.6 |
| Perineural invasion | Yes | 5 | 20.8 |
|  | No | 19 | 79.2 |
| Lymph node metastases | Yes | 10 | 41.7 |
|  | No | 14 | 58.3 |

The quantification of mRNA expression of HPSE, HPSE2, MMP-9, and SYND1 was obtained through quantitative RT-PCR analysis, and the values represent ΔCt relative expression of each marker and endogenous genes RPL13A and GAPDH, as shown in Fig 2.

**Fig 2.**
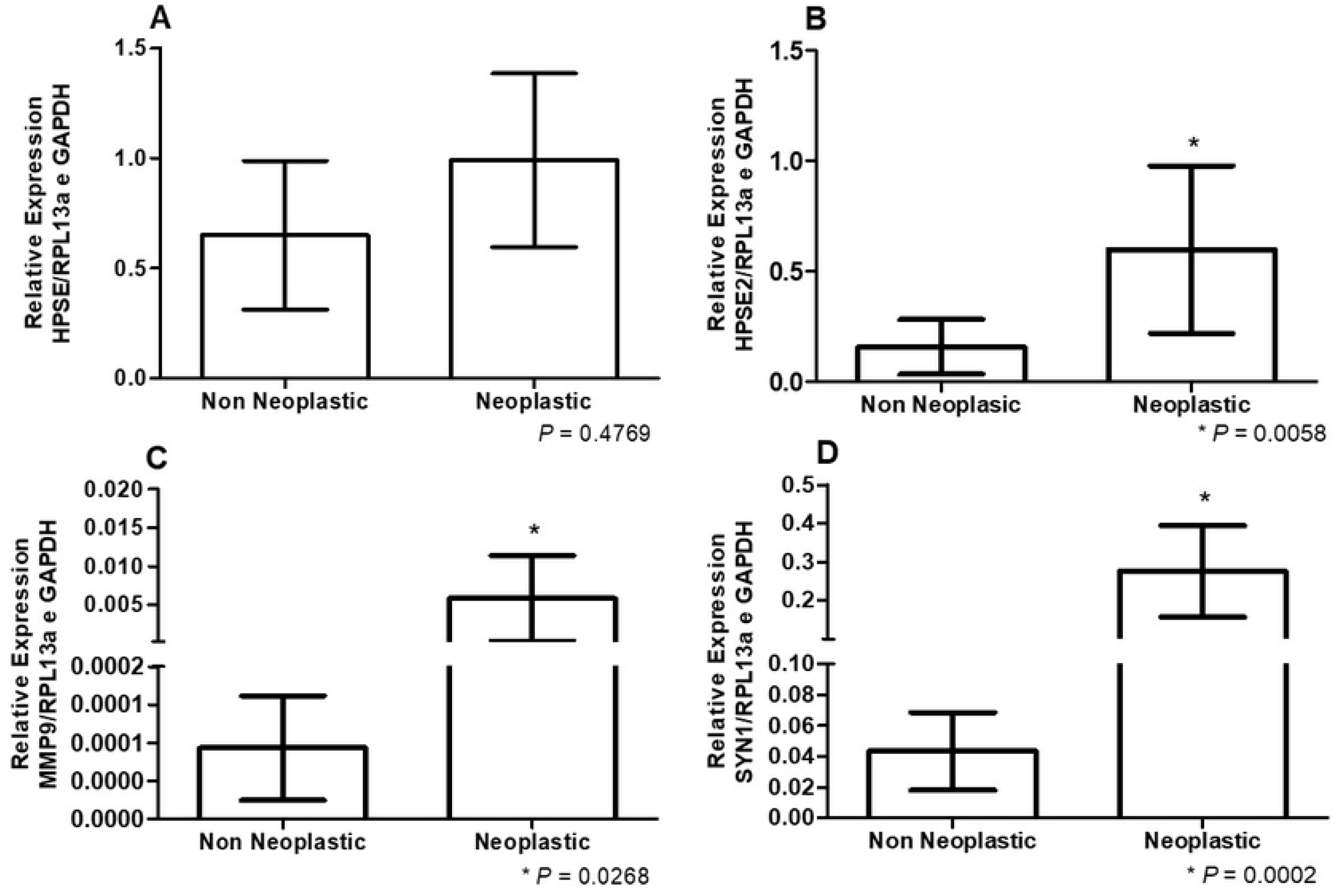
Comparative mRNA expression in nonneoplastic and neoplastic colorectal tissue. The results were obtained by quantitative RT-PCR performed on samples of neoplastic and nonneoplastic tissues of CRC patients. (A) heparanase expression (HPSE); (B) heparanase-2 expression (HPSE2); (C) metalloproteinase-9 expression (MMP-9) and (D) syndecan-1 expression (SYND1). The values represent the mean and standard deviation determined by 2^-ΔCt^ of triplicate assays with a total number of 24 samples in each group. The significance level (*P*) was obtained using the Student *t*-test.

We observed no difference regarding HPSE mRNA expression between neoplastic and nonneoplastic tissues obtained from CRC patients. However, HPSE2, MMP-9 and the heparan sulfate proteoglycan SYND1 have significantly increased mRNA expression in tumor tissues (Neoplastic) compared to the nonneoplastic samples (Fig 2).

There were significantly changes in the protein expression comparing tumor and nonaffected tissues from CRC patients. Immunohistochemistry analyzes clearly demonstrated higher protein level of HPSE, HPSE2, MMP-9 and SYND1 in neoplastic samples (Fig 3).

**Fig 3.**
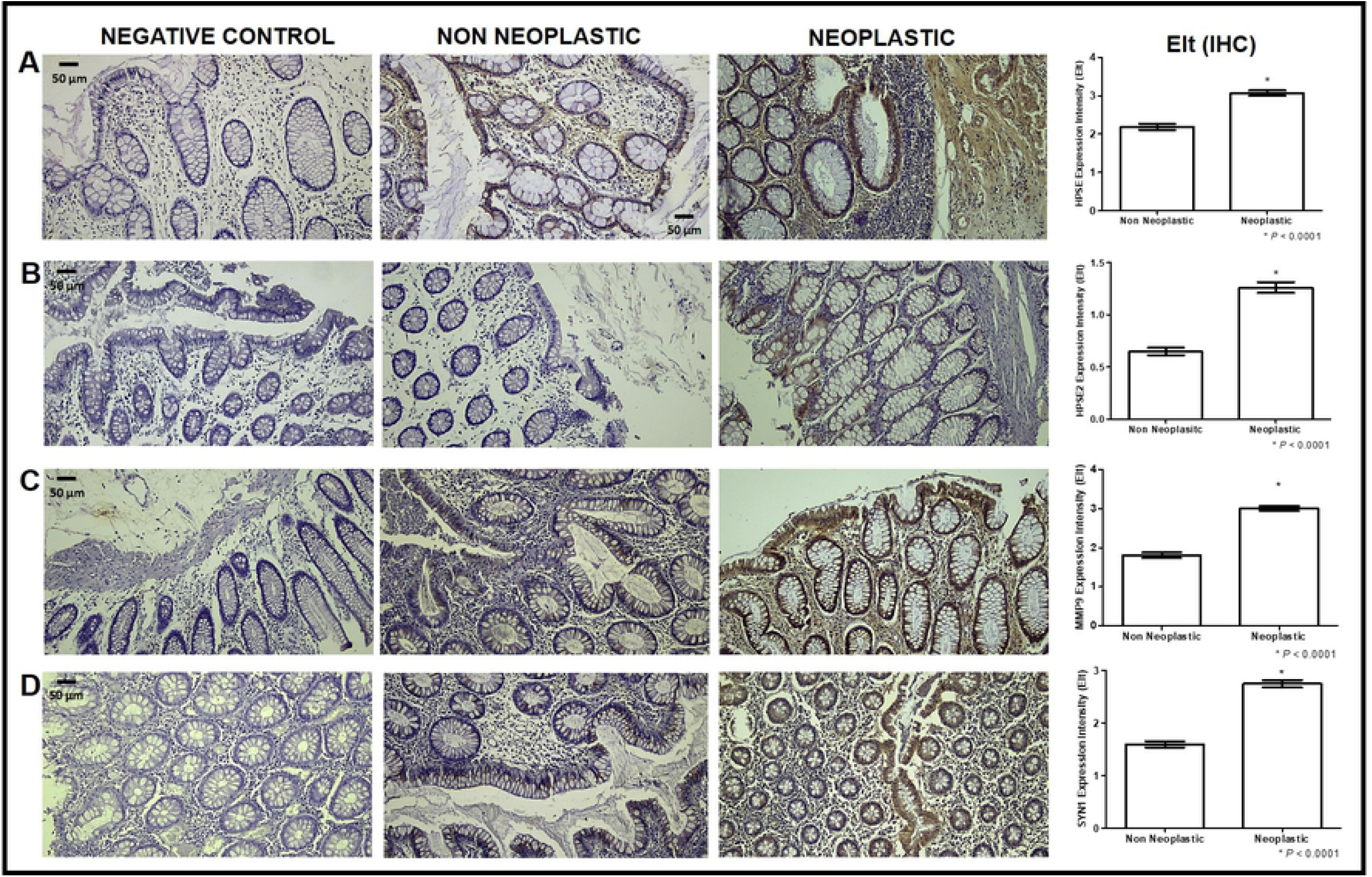
Comparative protein expression in nonneoplastic and neoplastic colorectal tissue. Expression intensity (Elt) was obtained after immunohistochemistry using specific antibodies. The negative control, absence of primary antibody. (A) HPSE, anti-heparanase-1 (HPA C-20); (B) HPSE2, anti-heparanase-2 (HPA C-17); (C) MMP-9, anti-metalloproteinase-9 (MMP-9 C-20) and (D), SYND1, anti-syndecan-1 (MCA681). Histomorphometry analysis was performed by two observers and one pathologist. A Nikon brand binocular microscope was used. Initially, an analysis of two dorsal quadrants was performed under 10X magnification and then the analysis was performed on 5-10 concentric fields under 40X magnification. The protein values were obtained using semiquantitative scoring as followed absent (zero); mild (1); moderate (2); intense (3), very intense (4). Chi-square test adopted a statistical significance level of 5% (*P* ≤ 0.05). N= 24 tissue samples.

We divided the patients in two groups, those free of lymph node metastasis (G1) and patients who had lymph node metastasis (G2) in order to investigate whether differences between HPSE, HPSE2, MMP-9, and SYND1 expression and lymph node metastasis occurrence. For this study, we evaluated the expression of the mRNA, since RT-PCR results represent a more precise quantitative analysis.

Fig 4 shows that only HPSE2 has a direct correlation with lymph node metastases, since significantly higher HPSE2 mRNA expression was observed in patients with lymph node metastases (*P* = 0.048).

**Fig 4.**
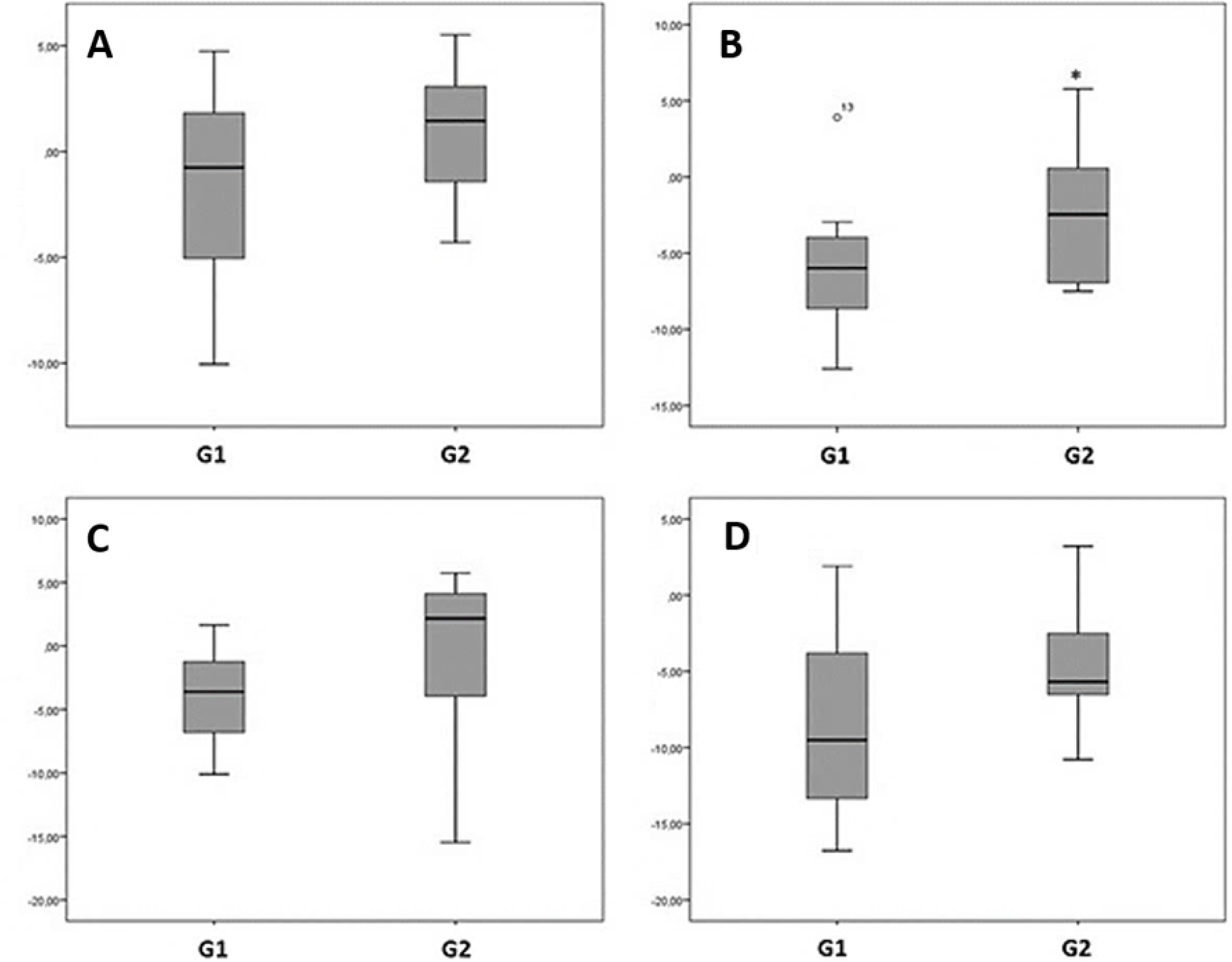
Comparative mRNA expression between patients without and with lymph node metastasis. G1, group of patients without lymph node metastasis (N=14) and G2, group of patients with lymph node metastasis (N=10). Relative expression of HPSE (A), HPSE2 (B), MMP-9 (C), and SYND1 (D). The analyses were performed by quantitative RT-PCR using mRNA expression of each marker in tumor samples (ΔCt), relative to the expression of nonneoplastic tissue (ΔΔCt). The values represent the medians and percentiles of assays performed in triplicate. The significance level (*P* = 0.048) was obtained using the Mann-Whitney test.

Table 2 represents the values of median and percentiles obtained by boxplot analysis of relative mRNA expression of HPSE, HPSE2, MMP-9 and SYND1 in patients with CRC without or with lymph node metastasis, respectively, Group 1 and Group 2.

**Table 2.**
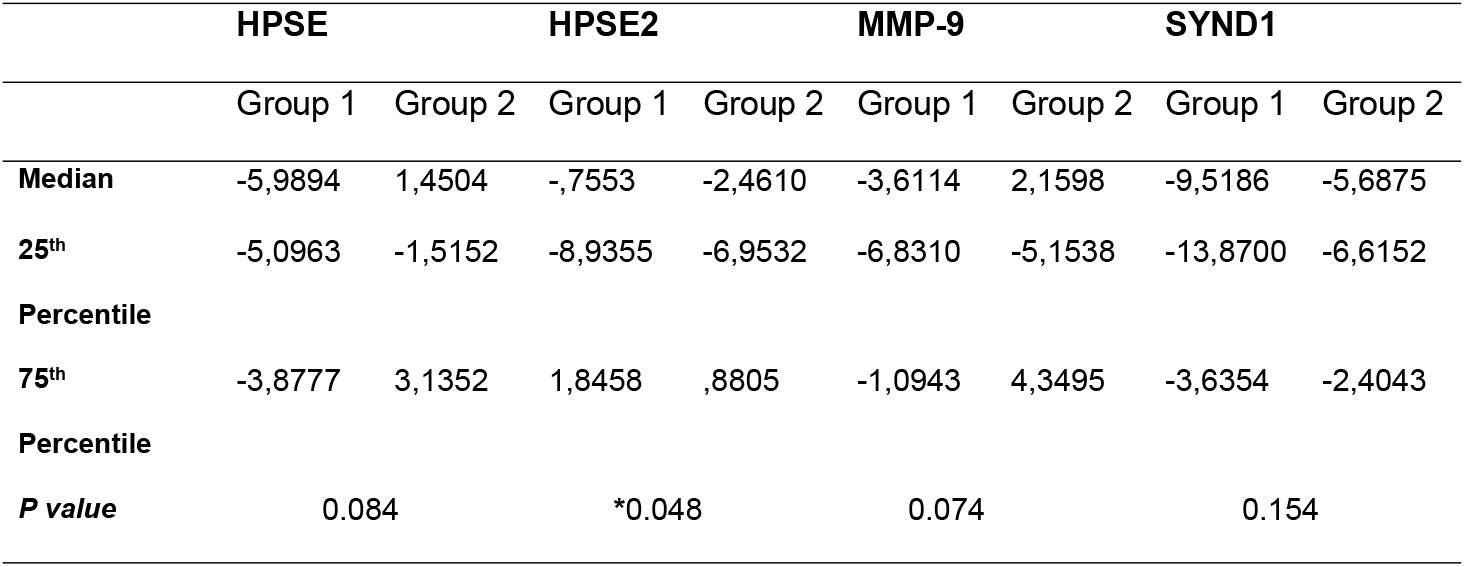
Relative expression of HSPE, HPSE2, MMP-9, and SYND1 in CRC Group 1 and Group 2, tissues of patients without and with lymph node metastasis, respectively.

## Discussion

The results of this study show that significant differences exist regarding the mRNA and protein expression of HPSE2, MMP-9, and SYND1 between neoplastic and non-neoplastic CRC tissues. The increased expression of such constituents has already been described in another study that compared tumor tissue with control tissue collected from individuals not affected by cancer [2].

Research has shown that HPSE is directly related to carcinogenesis. HPSE expression increased in cancer patients’ tumor tissues compared with tissues collected from individuals not affected by cancer. However, we decided to investigate whether the expression of HPSE differs between tumorous and non-tumorous tissues of a given cancer patient. The results showed no difference in the HPSE mRNA expression; however, protein expression demonstrated differences comparing tumor and adjacent nontumor tissues. The difference in the protein with no change in the mRNA expression of HPSE can be explained by a possible control of gene expression that occurs during protein synthesis, such as temporal regulation, and not in the transcription level. This result was highly relevant to our study, suggesting that the enzyme’s mRNA expression is already altered in tissue that is unaffected by the tumor.

Therefore, the present study’s data suggest that the HPSE enzyme might be a systemic marker, meaning that increased HPSE expression is not restricted to tumor tissue. Results obtained by our group showed an increase in the mRNA expression and protein level of HPSE in circulating lymphocytes of patients with breast cancer compared to women not affected by cancer [13], reinforcing the hypothesis that mRNA of HPSE is a possible biomarker that may not only be overexpressed in tumor cells.

Friedmann et al. did not detect the HSPE protein in the non-neoplastic colon epithelium adjacent to the CRC. In contrast, the HPSE gene and protein were already expressed at the colon adenoma stage, which was also observed in the present study [27]. In our previous study, the immunohistochemical analysis revealed a negative correlation between HPSE and HPSE2 proteins in colorectal adenomas [37]. However, the present results clearly showed an increased protein level of both HPSE and HPSE2 in CRC samples compared to adjacent tissues. Moreover, the enhanced HPSE2 protein expression detected in the present study corroborates our group’s previous study [2].

In this present study, the finding of increased HPSE2 expression in tumoral tissues in patients with lymph node metastases from CRC, compared to the HPSE2 expression in tumor tissue obtained from patients without lymph node metastases suggests that this protein may be related to molecular mechanisms of invasion and migration of tumor cells. Data from the literature have already proven a differential expression of HPSE2 in the cervix, endometrium, and thyroid cancer [38–40]. The fact that increased HPSE2 expression is directly related to tumors of patients with lymph node metastases possibly indicates that the HPSE2 expression might be useful to evaluate lymph node metastasis events. It could be a potential marker of worse prognosis for CRC patients [41]. Although the present study showed that there was an increase of HPSE2 in the group with lymph node metastasis, a larger N could allow subdivision of the samples in different stages of the disease and possibly point to a difference in the expression of HPSE in the group of patients with more advanced stages of cancer.

Day et al. used immunohistochemistry to study the changes in SYND1 expression during the development of colorectal neoplasia in adenomas and carcinomas arising from adenomas [5]. These results indicated the importance of cell-matrix adhesion disruption for colorectal carcinogenesis, anticipating changes in the cell-cell adhesion process intermediated by E-cadherin. The present investigation results indicated that HS proteoglycan SYND1 has higher protein and mRNA expression in tumor tissues than non-neoplastic tissues. Such a result was also observed in our group’s previous study, which found a significant decrease in CRC tissue’s SYND1 protein level than the non-neoplastic colorectal tissue [37]. It is important to note that such studies compared samples from tumor tissues and tissues of patients not affected by neoplasia. In the present study, we evaluated tumor and adjacent nonaffected tissues obtained from the same patient with CRC. We observed an increase in mRNA and protein of SYND1, which shows the care that must be taken to report the alterations of such proteoglycans in the carcinogenesis.

The association of SYND1 expression with pathological and prognostic aspects in patients with CRC produced controversial results. Wei et al. observed a correlation between loss of SYND1 expression and histological grade and CRC stage [3]. However, these authors did not identify a correlation between loss of SYND1 expression with the presence of lymph nodes compromised by the neoplasia or distant metastases. Fujiya et al. showed that patients with CRC and without expression of SYND1 had a higher incidence of lymph node and liver metastases, and survival was significantly lower regardless of the TNM stage [28]. Wang et al. reported that the lower disease-free survival in CRC was significantly associated with the elevated preoperative serum level of SYND1 [29]. Hashimoto et al. and Mitselou et al. did not observe that SYND1 expression in CRC affects patients’ prognosis with CRC [30,31].

On the other hand, Lundin et al. showed that 5-year survival in CRC patients did not change in patients with strong and weak SYND-1 expression [32]. In the present study, HS proteoglycan SYND1 exhibited higher expression of mRNA and protein in tumor tissues compared to the non-neoplastic tissues obtained from CRC patients. This finding may reinforce the idea that the development of HPSE inhibitor drugs to block HSPE/SYND1 axis activity and immunotherapy using anti-SYND1 antibodies may alleviate the aggressive character of malignant neoplasms [38,42].

In several malignant neoplasms, including CRC, increased levels of MMP-9 were observed during the carcinogenesis of these tumors, and this finding is associated with angiogenesis, tumor progression, invasion, metastasis, and lower patient survival rates [26,34,35]. Studies comparing the MMP-9 messenger RNA (mRNA) expression levels in the tissue from patients with CRC revealed increased expression compared to normal intestinal mucosa and adenomas [34]. Zhang et al.studied the MMP-9 levels in the tissues of patients with CRC and individuals without CRC [35]. The authors observed that MMP-9 levels were significantly elevated in patients with metastases. Similarly, they observed a 5-fold higher MMP-9 in CRC compared with non-neoplastic mucosa [34]. MMP-9 expression in CRC cells showed no significant relationship with the presence of CRC metastatic lymph nodes [35].

These data were similar to Langers et al. results; they also found no association between tissue level of MMP-9 and the presence of metastases in patients with CRC [26]. In contrast, MMP-9 up-regulation has also been described in CRC precursor lesions, which may indicate its use as a biomarker of early CRC diagnosis [22,36]. Some studies have presented conflicting results regarding the role of MMP-9 expression in the prognosis in CRC patients [33–36]. Immunohistochemical staining confirmed that MMP-9 is abundantly present in CRC cells and that tumor-associated macrophages are significant sources of MMP in the carcinogenesis process [35]. This result corroborates with increased mRNA and protein expression of MMP-9 in tumor samples obtained in the present study. The enhanced MMP-9 expression was correlated with poor outcome and suggested an attractive therapeutic target [26,33].

Our study was limited by its retrospective nature and the selection criteria. The cohort contains a relatively small population of patients with CRC but without preoperative treatment (chemotherapy or radiotherapy).

The combined analysis of the expression of mRNA and protein significantly strengthen the findings.

In conclusion, both mRNA and protein expression of HPSE2, MMP-9, and SYND1 are increased in tumor samples than the adjacent tissue collected from patients with CRC, indicating that such molecules might be involved with ECM changes that occur during carcinogenesis. The increase in HPSE protein level without altering mRNA expression may suggest that the control of gene expression of such enzyme possibly occurs at the level of protein synthesis rather than transcription. The increased HPSE2 expression in patients with lymph node metastasis possibly indicates that HPSE2 can participate in this event in colorectal carcinoma.

## Author contributions statement

T.R.T., K.C.T., R.L.S., J.W. and M.A.S.P. performed data analysis. T.R.T. and M.A.S.P. designed the study. R.L.S., M.A.F.R.J. and J.W. contributed to acquisition of data. T.R.T., K.C.T., S.S.S. and J.W., did literature review. T.R.T., K.C.T., S.S.S., M.A.F.R.J., J.W. and M.A.S.P. drafted the manuscript. S.S.S., R.L.S., M.A.F.R.J. and J.W. did medical and technical analysis. M.A.S.P. revised the manuscript. All authors reviewed the manuscript.

## Additional information

The authors declare no competing interests.

All data generated or analyzed during this study are included in this published article (and its Supplementary Information files).

## References

1. Arnold M, Sierra MS, Laversanne M, Soerjomataram I, Jemal A and Bray F. Global patterns and trends in colorectal cancer incidence and mortality. Gut. 2017;66: 683–691. http://dx.doi.org/10.1136/gutjnl-2015-310912.

2. Peretti T, Waisberg J, Mader AMAA, de Matos LL, da Costa RB, Conceição GM, et al. Heparanase-2, syndecan-1, and extracellular matrix remodeling in colorectal carcinoma. Eur. J. Gastroenterol. Hepatol. 2008;20: 756–765. https://doi.org/10.1097/MEG.0b013e3282fc2649.

3. Wei HT, Guo EN, Dong BG and Chen LS. Prognostic and clinical significance of syndecan-1 in colorectal cancer: a meta-analysis. BMC Gastroenterol. 2015;15: 152. https://doi.org/10.1186/s12876-015-0383-2.

4. Shteingauz A, Ilan N and Vlodavsky I. Processing of heparanase is mediated by syndecan-1 cytoplasmic domain and involves syntenin and α-actinin. Cell Mol Life Sci. 2014;71: 4457–4470. https://doi.org/10.1007/s00018-014-1629-9.

5. Day RM, Hao X, Ilyas M, Daszak P, Talbot IC and Forbes A. Changes in the expression of syndecan-1 in the colorectal adenoma-carcinoma sequence. Virchows Arch. 1999;434: 121–125. https://doi.org/10.1007/s004280050315.

6. Qing Q, Zhang S, Chen Y, Li R, Mao H and Chen Q. High glucose-induced intestinal epithelial barrier damage is aggravated by syndecan-1 destruction and heparanase overexpression. J Cell Mol Med. 2015;19: 1366–1374. https://doi.org/10.1111/jcmm.12523.

7. Rodrigues LM, Theodoro TR, de Matos LL, Mader AMAA, Milani C and Pinhal MAS. Heparanase isoform expression and extracellular matrix remodeling in intervertebral disc degenerative disease. Clinics. 2011;66: 903–909. http://dx.doi.org/10.1590/S1807-59322011000500030.

8. Shteingauz A, Boyango I, Naroditsky I, Hammond E, Gruber M, Doweck I, et al. Heparanase Enhances Tumor Growth and Chemoresistance by Promoting Autophagy. Cancer Res. 2015;75: 3946–3957. http://doi.org/10.1158/0008-5472.CAN-15-0037.

9. Sanderson RD, Elkin M, Rapraeger AC, Ilan N, Vlodavsky I. Heparanase regulation of cancer, autophagy and inflammation: New mechanisms and targets for therapy. FEBS J. 2017; 284: 42–55. https://doi.org/10.1111/febs.13932.

10. Levy-Adam F, Feld S, Cohen-Kaplan V, Shteingauz A, Gross M, Arvatz G, et al. Heparanase 2 interacts with heparan sulfate with high affinity and inhibits heparanase activity. J Biol Chem. 2010;285: 28010–28019. https://doi.org/10.1074/jbc.M110.116384.

11. Vlodavsky I, Singh P, Boyango I, Gutter-Kapon L, Elkin M, Sanderson RD, et al. Heparanase: From basic research to therapeutic applications in cancer and inflammation. Drug Resist Updat. 2016;29: 54–75. https://doi.org/10.1016/j.drup.2016.10.001.

12. Dempsey LA, Brunn GJ and Platt JL. Heparanase, a potential regulator of cell-matrix interactions. Trends Biochem Sci. 2000;25: 349–351. https://doi.org/10.1016/S0968-0004(00)01619-4.

13. Theodoro TR, de Matos LL, Anna AVLS, Fonseca FLA, Semedo P, Martins LC, et al. Heparanase expression in circulating lymphocytes of breast cancer patients depends on the presence of the primary tumor and/or systemic metastasis. Neoplasia. 2007;9: 504–510. https://doi.org/10.1593/neo.07241.

14. Yang Y, MacLeod V, Miao HQ, Theus A, Zhan F, Shaughnessy JD, et al. Heparanase enhances syndecan-1 shedding: a novel mechanism for stimulation of tumor growth and metastasis. J Biol Chem. 2007;282: 13326–13333. https://doi.org/10.1074/jbc.M611259200.

15. Masola V, Secchi MF, Gambaro, G and Onisto M. Heparanase as a target in cancer therapy. Curr Cancer Drug Targets. 2014;14: 286–293. https://doi.org/10.2174/1568009614666140224155124.

16. Sato T, Yamaguchi A, Goi T, Hirono Y, Takeuchi K, Katayama K., et al. Heparanase expression in human colorectal cancer and its relationship to tumorangiogenesis, hematogenous metastasis, and prognosis. J Surg Oncol. 2004;87: 174–181. https://doi.org/10.1002/jso.20097.

17. Edovitsky E, Elkin M, Zcharia E, Peretz T and Vlodavsky I. Heparanase gene silencing, tumor invasiveness, angiogenesis, and metastasis. J Natl Cancer Ins. 2004;96: 1219–1230. https://doi.org/10.1093/jnci/djh230.

18. Lerner I, Hermano E, Zcharia E, Rodkin D, Bulvik R, Doviner V, et al. Heparanase powers a chronic inflammatory circuit that promotes colitis-associated tumorigenesis in mice. J Clin Invest. 2011;121: 1709–1721. https://doi.org/10.1172/jci43792.

19. Theodoro TR, Matos LL, Cavalheiro RP, Justo GZ, Nader HB and Pinhal MAS. Crosstalk between tumor cells and lymphocytes modulates heparanase expression. J Transl Med. 2019;17: 103. https://doi.org/10.1186/s12967-019-1853-z.

20. Singh P, Blatt A, Feld S, Zohar Y, Saadi E, Barki-Harrington L, et al. The Heparanase Inhibitor PG545 Attenuates Colon Cancer Initiation and Growth, Associating with Increased p21 Expression. Neoplasia. 2017;19: 175–184. https://doi.org/10.1016/j.neo.2016.12.001.

21. Zarkavelis G, Boussios S, Papadaki A, Katsanos KH, Christodoulou DK and Pentheroudakis G. Current and future biomarkers in colorectal cancer. Ann Gastroenterol. 2017;30: 613–621. https://doi.org/10.20524/aog.2017.0191.

22. Jonsson A, Hjalmarsson C, Falk P and Ivarsson ML. Stability of matrix metalloproteinase-9 as biological marker in colorectal cancer. Med Oncol. 2018;35: 50. https://doi.org/10.1007/s12032-018-1109-4.

23. Herszenyi L, Hritz I, Lakatos G, Varga MZ and Tulassay Z. The behavior of matrix metalloproteinases and their inhibitors in colorectal cancer. Int J Mol Sci. 2012;13: 13240–13263. https://doi.org/10.3390/ijms131013240.

24. Marshall DC, Lyman SK, McCauley S, Kovalenko M, Spangler R, Liu C, et al. Selective allosteric inhibition of MMP-9 is efficacious in preclinical models of ulcerative colitis and colorectal cancer. PLoS One. 2015;10: 5. https://doi.org/10.1371/journal.pone.0127063.

25. Bendardaf R, Buhmeida A, Hilska M, Laato M, Syrjänen S, Syrjänen K, et al. MMP-9 (gelatinase B) expression is associated with disease-free survival and disease-specific survival in colorectal cancer patients. Cancer Invest. 2010;28: 38–43. https://doi.org/10.3109/07357900802672761.

26. Langers AMJ, Verspaget HW, Hawinkels LJAC, Kubben FJGM, Van Duijn W, Van Der Reijden JJ, et al. MMP-2 and MMP-9 in normal mucosa are independently associated with outcome of colorectal cancer patients. Br J Cancer. 2012;106: 1495–1498. https://doi.org/10.1038/bjc.2012.80.

27. Friedmann Y, Vlodavsky I, Aingorn H, Aviv A, Peretz T, Pecker I, et al. Expression of Heparanase in Normal, Dysplastic, and Neoplastic Human Colonic Mucosa and Stroma: Evidence for Its Role in Colonic Tumorigenesis. Am J Pathol. 2000;157: 1167–1175. https://doi.org/10.1016/S0002-9440(10)64632-9.

28. Fujiya M, Watari J, Ashida T, Honda M, Tanabe H, Fujiki T, et al. Reduced expression of syndecan-1 affects metastatic potential and clinical outcome in patients with colorectal cancer. Jpn J Cancer Res. 2001;92: 1074–1081. https://doi.org/10.1111/j.1349-7006.2001.tb01062.x.

29. Wang X, Zuo D, Chen Y, Li W, Liu R, He Y, et al. Shed Syndecan-1 is involved in chemotherapy resistance via the EGFR pathway in colorectal cancer. Br J Cancer. 2014;111: 1965–1976. https://doi.org/10.1038/bjc.2014.493.

30. Hashimoto Y, Skacel M and Adams JC. Association of loss of epithelial syndecan-1 with stage and local metastasis of colorectal adenocarcinomas: an immunohistochemical study of clinically annotated tumors. BMC Cancer. 2008;8: 185. https://doi.org/10.1186/1471-2407-8-185.

31. Mitselou A, Skoufi U, Tsimogiannis KE, Briasoulis E, Vougiouklakis T, Arvanitis D, et al. Association of syndecan-1 with angiogenesis-related markers, extracellular matrix components, and clinicopathological features in colorectal carcinoma. Anticancer Res. 2012;32: 3977–3985. PMID: 22993347.

32. Lundin M, Nordling S, Lundin J, Isola J, Wiksten JP and Haglund C. Epithelial syndecan-1 expression is associated with stage and grade in colorectal cancer. Oncology. 2005;68: 306–313. https://doi.org/10.1159/000086969.

33. Said AH, Raufman JP and Xie G. The role of matrix metalloproteinases in colorectal cancer. Cancer. 2014;6: 366–375. https://doi.org/10.3390/cancers6010366.

34. Hoelzle CR, Magalhães KC, Carvalho SS, Santos GA, Maia IM, Sousa MC, et al. Matrix metalloproteinase 9-1562C/T polymorphism increased protein levels in patients with colorectal cancer in a sample from southeastern Brazil. Genet Mol Res. 2016;15: 1. https://doi.org/10.4238/gmr.15017478.

35. Yi Zhang, Xiao-Ya Guan, Bin Dong, Min Zhao, Jian-Hui Wu, Xiu-Yun Tian, et al. Expression of MMP-9 and WAVE3 in colorectal cancer and its relationship to clinicopathological features. J Cancer Res Clin Oncol. 2012;138: 2035–2044. https://doi.org/10.1007/s00432-012-1274-3.

36. Otero-Estévez O, De Chiara L, Rodríguez-Girondo M, Rodríguez-Berrocal FJ., Cubiella J, Castro I, et al. Serum matrix metalloproteinase-9 in colorectal cancer family-risk population screening. Sci Rep. 2015;5: 13030. https://doi.org/10.1038/srep13030.

37. Waisberg J, Theodoro TR, Matos LL, Brasil F, Serrano RL, Saba GT and Pinhal MAS. Immunohistochemical expression of heparanase isoforms and syndecan-1 proteins in colorectal adenomas. Eur J Histochem. 2016;60: 2590. https://doi.org/10.4081/ejh.2016.2590.

38. Marques RM, Focchi GR, Theodoro TR, Castelo A, Pinhal MA and Nicolau SM. The immunoexpression of heparanase 2 in normal epithelium, intraepithelial, and invasive squamous neoplasia of the cervix. J Low Genit Tract Dis. 2012;16: 256–262. https://doi.org/10.1097/LGT.0b013e3182422c69.

39. Matos LL, Suarez ER, Theodoro TR, Trufelli DC, Melo CM, Garcia LF, et al. The profile of heparanase expression distinguishes differentiated thyroid carcinoma from benign neoplasms. PLoS One. 2015;10: e0141139. https://doi.org/10.1371/journal.pone.0141139.

40. Signorini Filho RC, de Azevedo Focchi GR, Theodoro TR, Pinhal MA and Nicolau SM. Immunohistochemical expression of heparanases 1 and 2 in benign tissue and in invasive neoplasia of the endometrium: a case-control study. Int J Ginecol Cancer. 2015;25: 269–278. http://dx.doi.org/10.1097/IGC.0000000000000329.

41. Thompson CA, Purushothaman A, Ramani VC, Vlodavsky I and Sanderson RD. Heparanase regulates secretion, composition, and function of tumor cell-derived exosomes. J Biol Chem. 2013;288: 10093–10099. http://dx.doi.org/10.1074/jbc.C112.444562.

42. Thompson CA, Purushothaman A, Ramani VC, Vlodavsky I and Sanderson RD. The heparanase/syndecan-1 axis in cancer: mechanisms and therapies. FEBS J. 2013;280: 2294–2306. https://doi.org/10.1111/febs.12168.

